# Pre-myelinating oligodendrocyte ADGRG1 is required for axon ensheathment and CNS myelin formation

**DOI:** 10.64898/2026.01.26.701898

**Authors:** Beika Zhu, Tao Li, Benjamin Rodrigues, Melanie Piller, Cody L. Call, Amit Mogha, Arantzazu Belloso Iguerategui, Brian Chiou, Rachael Schmidt, Andi Wangzhou, Alicia L. Thurber, Madison Dee, Claire H. Larson, Kelly R. Monk, Xianhua Piao

## Abstract

Myelin is essential for axonal health and rapid propagation of action potentials. In the central nervous system (CNS), myelination is initiated with axon ensheathment by pre-myelinating oligodendrocytes (preOLs), followed by iterative membrane wrapping and longitudinal extension along the axon. The molecular mechanisms that govern preOL development remain largely unknown. In this study, we identify the adhesion G protein–coupled receptor ADGRG1 (also called GPR56) as a key, evolutionarily conserved regulator of this process. We show that ADGRG1 is highly expressed in preOLs and that its conditional deletion in mouse preOLs leads to simplified preOL morphology and defective axon ensheathment, resulting in CNS hypomyelination. Live imaging in zebrafish demonstrated conserved function of Adgrg1 in axon ensheathment. Mechanistically, we show that ADGRG1 promotes preOL maturation by RhoA activation. Collectively, these findings reveal an ADGRG1-mediated RhoA signaling pathway that governs axon ensheathment and myelin formation.

## Introduction

Myelin, a specialized multilamellar membrane structure that wraps around axons, is critical for axonal health and efficient nerve conduction (*1, 2*). Disruption of myelin leads to a broad spectrum of neurological disorders, including multiple sclerosis (*3, 4*), leukodystrophies (*5*), and various neurodegenerative diseases (*6, 7*). Central nervous system (CNS) myelin is produced by oligodendrocytes (OLs). OL development follows a well-defined developmental lineage progression from proliferating oligodendrocyte precursor cells (OPCs) to pre-myelinating oligodendrocytes (preOLs) and finally to myelinating oligodendrocytes (mOLs) (*8*). The preOL stage serves as a critical transition point where OLs undergo extensive morphological transformations, extending dynamic processes to seek out and initiate the wrapping of axons (*9, 10*). Defining the molecular regulators that govern this specific developmental transition stage is crucial for understanding myelination and for developing strategies to promote myelin repair.

Several extrinsic cues and intracellular signaling pathways have been implicated in myelination (*8*), such as integrin-ECM interactions and focal adhesion kinase (FAK) signaling (*11, 12*), Notch signaling (*13, 14*), the Wnt-β-catenin (*15, 16*), and the RhoA signaling pathway (*17–19*). However, the identity of upstream transmembrane receptors that coordinate these signals and precisely regulate preOL timely maturation and initiation of axon ensheathment remains largely unknown.

G protein-coupled receptors (GPCRs) represent a large family of transmembrane receptors with diverse roles in neural development. How GPCRs function in preOLs during axon ensheathment is largely unexplored. In this study, we mined a published single-cell RNA sequencing dataset (*20*) to profile GPCR expression in preOLs. We discovered that the adhesion GPCR ADGRG1 (also known as GPR56) is highly expressed in preOLs. ADGRG1 has been implicated in myelination—deleting *Adgrg1* in OPCs disrupts oligodendrocyte proliferation and leads to myelination deficits (*18, 21, 22*). Here, we demonstrate a unique, indispensable, and conserved role of ADGRG1 in preOL development and function in mice and zebrafish. Targeted deletion of *Adgrg1* in preOLs impairs their maturation and disrupts axon ensheathment, leading to CNS hypomyelination. Mechanistically, we find that ADGRG1 signals through the RhoA pathway to drive this process. By identifying ADGRG1 as a key intrinsic regulator of OL maturation and axon ensheathment, we provide novel insights into the control of myelination and identify potential therapeutic targets for enhancing remyelination.

## Results

### ADGRG1 is highly expressed in pre-myelinating oligodendrocytes

To identify molecular regulators of preOLs, we took advantage of a published single-cell RNA sequencing (scRNAseq) dataset (*20*) and performed unbiased clustering. We identified three distinct OL lineage clusters: OPCs, preOLs, and myelinating oligodendrocytes (mOLs), in line with previous studies (*20, 23*) **(Figure 1A)**. Given the established roles of GPCRs in neural development, we examined their expression patterns across the preOL and mOL clusters. Differential expression analysis revealed 13 GPCRs with significantly altered expression levels, 5 downregulated and 8 upregulated, in preOLs **(Figure 1B)**. Among the upregulated receptors, *Adgrg1* stood out due to its robust and selective enrichment in preOLs **(Figures 1B-C)**. In contrast, the expression of *Plp1* was initiated in the preOL stage and sustained in mOLs, but largely absent in OPCs **(Figure 1D)**.

**Figure 1.**
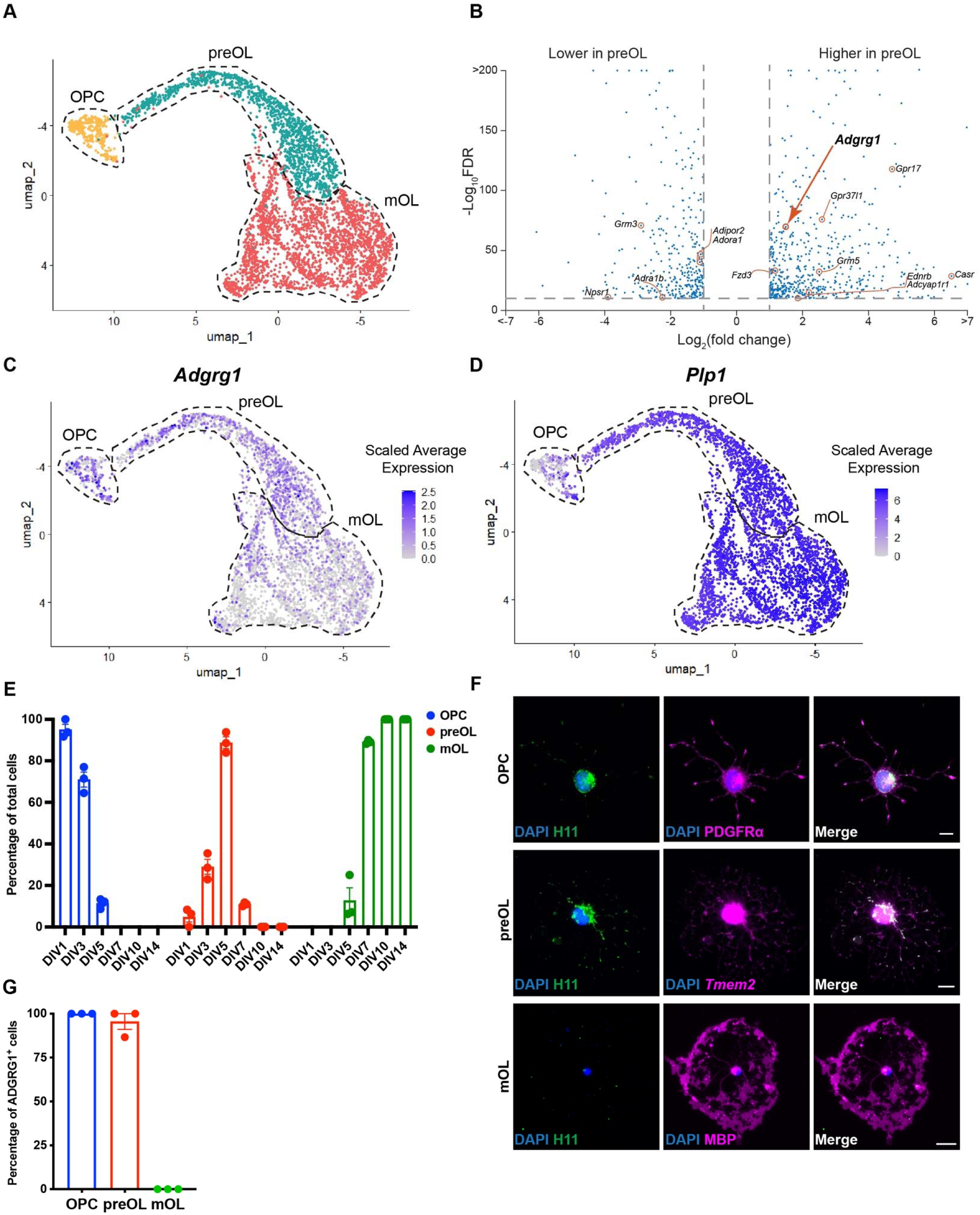
*Adgrg1* is highly expressed in pre-myelinating oligodendrocytes. **(A)** UMAP of single cell RNAseq showing oligodendrocyte lineage (OL) progression from OL precursor cells (OPCs) to pre-myelinating OL (preOLs) and mature OL (mOLs). **(B)** Volcano plot of differentially expressed genes between preOLs and mOLs. All genes encoding GPCRs with > 2-fold changes and < 10^-10^ FDR are highlighted. **(C-D)** UMAP feature plots showing expression of *Adgrg1* **(C)** and *Plp1* **(D)** across the oligodendrocyte lineage. **(E)** Quantification of the lineage progression in culture over a 14-day time course, showing the percentage of OPCs, preOLs, and mOLs at different differentiation time course. Each circle represents one biological replicate. **(F)** Representative immunocytochemistry of primary mouse OLs at different developmental stages. Cells were stained for ADGRG1 (H11, green), stage-specific markers (magenta) including PDGFRl7l for OPCs, *Tmem2* for preOLs, MBP for mOLs, and DAPI (blue). Scale bar, 10 µm. **(G)** Quantification showing the percentage of ADGRG1^+^ cells within the OPCs, preOLs and mOLs populations. Each circle represents one biological replicate. Data are presented as means ± SEM.

To validate these transcriptomic findings at the protein level, we used an established *in vitro* model in which immunopanned OPCs differentiate into preOLs and subsequently mOLs (*24*). This culture system successfully recapitulated the normal progression of oligodendrocyte maturation, with the proportion of OPCs decreasing over time while the proportions of preOLs and subsequently mOLs increased **(Figure 1E)**. We further demonstrated that ADGRG1 protein follows similar expression dynamics, with high protein levels in PDGFRα^+^ OPCs and *Tmem2*^+^ preOLs, while the signal was largely absent in MBP^+^ mOLs **(Figure 1F)**. Furthermore, almost all OPCs and preOLs were ADGRG1-positive, but ADGRG1 expression was mostly undetectable in mOLs **(Figure 1G)**. Together, these results indicate that ADGRG1 is selectively enriched in preOLs, supporting a potentially important role for this GPCR in the transition from precursor to myelinating OLs.

### ADGRG1 is required for the maturation of pre-myelinating oligodendrocytes

*Plp* is largely expressed during preOL and mOL stages whereas *Adgrg1* is mostly absent in mOLs **(Figures 1C and D)**. To investigate the function of ADGRG1 in preOLs, we utilized the *Plp-CreER* mouse line, a tool commonly used to label and manipulate genes starting from the preOL stage (*25*). We crossed *Adgrg1* floxed mice with *Plp-CreERT* mice to generate an inducible *Adgrg1* conditional knockout (*Adgrg1^fl/fl^; Plp^CreERT/+^*; cKO) and corresponding controls (*Adgrg1^+/+^; Plp^CreERT/+^*). To validate this model, we immunopanned OPCs from control and cKO mice, and confirmed that 4-hydroxy-tamoxifen (4OHT) treatment efficiently deleted ADGRG1 in preOLs but not in OPCs **(Figures S1A-B)**. Using this model, we first asked if preOL ADGRG1 regulates OL differentiation *in vitro*. Compared to controls, *Adgrg1* cKO cultures exhibited a significant and persistent delay in the generation of BCAS1 and MBP double positive OLs across a seven-day time course **(Figures 2A-C)**. To determine if this maturation defect occurred *in vivo*, we administered tamoxifen to pups from postnatal day 4 (P4) to P7, with booster doses on P10 and P11, to induce recombination during the preOL stage (*9*) **(Figure S2A)**. Using the Ai14 reporter, we observed that approximately 41% of TCF7L2^+^ (*26*) preOLs in the corpus callosum were successfully recombined **(Figures S2B-C)**. Analysis of the corpus callosum at P14 revealed a phenotype consistent with our *in vitro* findings. Compared to controls, cKO mice exhibited a sharp reduction in the number of BCAS1^+^ MBP^+^ OLs **(Figure 2D-E)**. This decrease was accompanied by a significant accumulation of aberrant BCAS1^+^ preOLs with disorganized and centrally distributed MBP signals **(Figure 2D and 2F)**. Taken together, these *in vitro* and *in vivo* data demonstrate that ADGRG1 is critically required for the maturation of preOLs.

**Figure 2.**
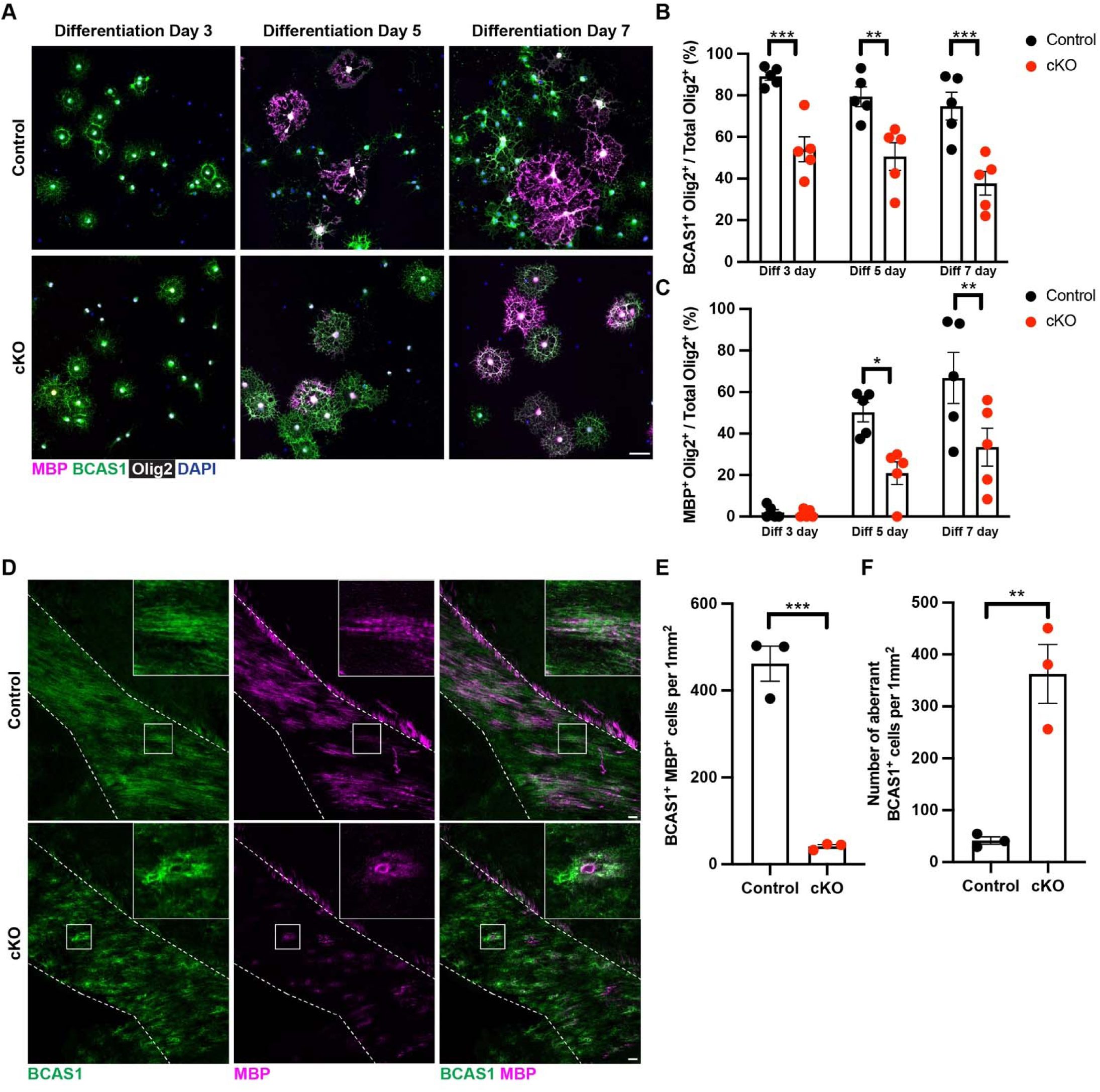
ADGRG1 promotes oligodendrocyte maturation. **(A)** Representative images of control and cKO oligodendrocyte cultures differentiated for 3, 5, or 7 days. Cultures were stained for MBP (magenta), BCAS1 (green), Olig2 (gray), and DAPI (blue). Scale bar, 10 μm. **(B)** Quantification of the percentage of BCAS1^+^ Olig2^+^ cells in Olig2^+^ population across differentiation time points. Each circle represents one biological replicate. *** p=0.0005 (for diff 3 day), ** p=0.0035 (for diff 5 day), *** p=0.0002 (for diff 7 day). Two-way ANOVA with Bonferroni’s multiple comparisons test. **(C)** Quantification of the percentage of MBP^+^ Olig2^+^ cells in Olig2^+^ population across differentiation time points. Each circle represents one biological replicate. * p=0.019, ** p=0.0072. Two-way ANOVA with Bonferroni’s multiple comparisons test. **(D)** Immunostaining of the corpus callosum for BCAS1 (green) and MBP (magenta) in tamoxifen injected P14 control and cKO mice. Dotted lines mark the corpus callosum boundary. Scale bar, 50 µm. Boxed areas are shown at higher magnification. **(E)** Quantification of the percentage of BCAS1 and MBP double positive cells per 1mm^2^ in the corpus callosum. Each circle represents one biological replicate. *** p=0.0005. Unpaired two-tailed t-test. **(F)** Quantification of aberrant BCAS1^+^ cells per 1 mm^2^ in the corpus callosum. Each circle represents one biological replicate. ** p=0.0050. Unpaired two-tailed t-test. Data are presented as means ± SEM.

### ADGRG1 promotes preOL maturation through RhoA activation

Oligodendrocyte differentiation is accompanied by morphological transformation driven by the extension and stabilization of cellular processes. The RhoA signaling pathway is a primary intracellular regulator that governs these critical changes in cell shape and process dynamics (*27, 28*). Based on a prior study showing ADGRG1 can couple with Gα_12/13_ to activate the small GTPase RhoA (*18*), we hypothesized that ADGRG1 drives maturation through the RhoA signaling pathway in preOLs **(Figure 3A)**. To test this, we first examined if ADGRG1 activates RhoA in preOLs. Immunostaining revealed that in control preOLs, ADGRG1 colocalized with GTP-bound (active) RhoA, particularly in the processes. In contrast, GTP-RhoA signal was nearly undetectable in *Adgrg1* cKO preOLs, confirming that ADGRG1 activates RhoA in these cells **(Figure 3B)**.

**Figure 3.**
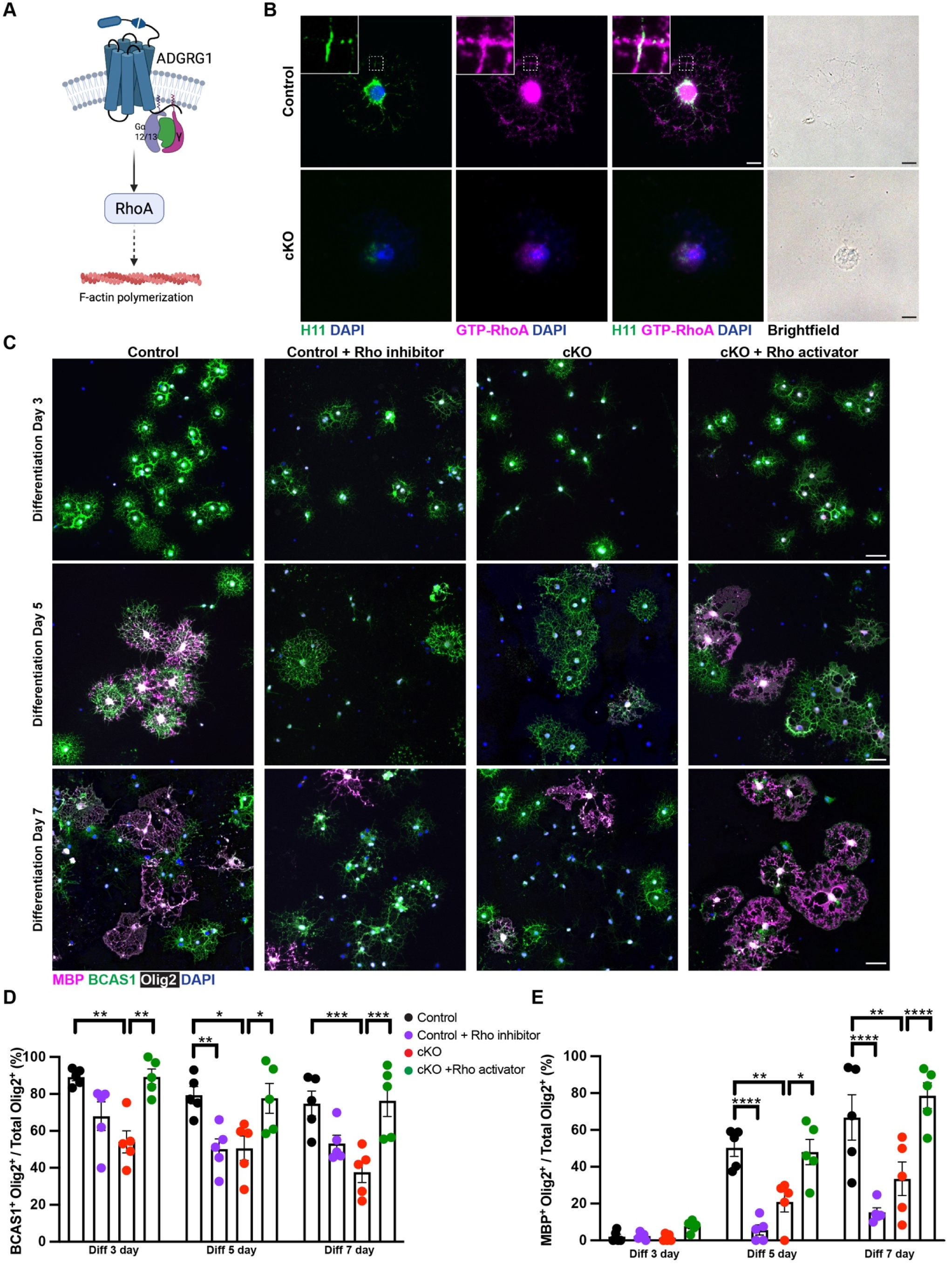
preOL *ADGRG1* regulates oligodendrocyte differentiation through RhoA pathway. **(A)** Schematic of the proposed signaling pathway showing ADGRG1 activating the RhoA pathway to promote F-actin polymerization in preOLs. **(B)** Immunostaining of control and cKO preOLs for ADGRG1 (H11, green), GTP-RhoA (magenta), DAPI (blue) and brightfield. Colocalization between ADGRG1 and GTP-RhoA was observed in the processes of control preOLs, but both signals were downregulated in cKO preOLs. Scale bar, 10 μm. **(C)** Representative images of control and cKO oligodendrocyte cultures differentiated for 3, 5, or 7 days, treated with either vehicle, Rho inhibitor (for control), or Rho activator (for cKO). Cultures were stained for MBP (magenta), BCAS1 (green), Olig2 (gray), and DAPI (blue). Scale bar, 10 μm. **(D)** Quantification of the percentage of BCAS1^+^ Olig2^+^ cells in total Olig2^+^ cells across differentiation time points. Each circle represents one biological replicate. ** p=0.0011 (diff 3 day Con vs cKO), ** p=0.0011 (diff 3 day cKO vs cKO + Rho activator), ** p=0.0088 (diff 5 day Con vs Con + Rho inhibitor), * p=0.0103 (diff 5 day Con vs cKO), * p=0.0181 (diff 5 day cKO vs cKO + Rho activator), *** p=0.0005 (diff 7 day Con vs cKO), *** p=0.0003 (diff 7 day cKO vs cKO + Rho activator). Two-way ANOVA with Bonferroni’s multiple comparisons test. **(E)** Quantification of the percentage of MBP^+^ Olig2^+^ cells in total Olig2^+^ cells across differentiation time points. Each circle represents one biological replicate. **** p<0.0001 (diff 5 day Con vs Con + Rho inhibitor), ** p=0.0047 (diff 5 day Con vs cKO), * p=0.0108 (diff 5 day cKO vs cKO + Rho activator), **** p<0.0001 (diff 7 day Con vs Con + Rho inhibitor), ** p=0.0010 (diff 7 day Con vs cKO), **** p<0.0001 (diff 7 day cKO vs cKO + Rho activator). Two-way ANOVA with Bonferroni’s multiple comparisons test. Data are presented as means ± SEM.

We next performed pharmacological treatments *in vitro* to determine if RhoA activity is both necessary for preOL differentiation and sufficient to rescue the phenotype associated with preOL *Adgrg1* deletion. Notably, inhibiting RhoA activity in controls phenocopied the *Adgrg1* cKO, significantly suppressing the generation of both BCAS1^+^ and MBP^+^ oligodendrocytes. Conversely, pharmacologically activating RhoA in *Adgrg1* cKOs was sufficient to rescue the maturation defects, restoring the number of BCAS1^+^ and MBP^+^ oligodendrocytes to control levels **(Figures 3C-E)**. Together, these results demonstrate that ADGRG1 promotes the timely maturation of preOLs by activating a RhoA-dependent signaling pathway. Pharmacological activation of RhoA is sufficient to bypass the requirement for upstream ADGRG1 signaling in driving oligodendrocyte differentiation.

### ADGRG1 is required for axon ensheathment and myelin formation

To determine the functional consequences of this maturation defect on myelin sheath formation, we first employed an *in vitro* microfiber assay to model axon ensheathment. Control and cKO OPCs were seeded on electrospun microfibers and differentiated for 14 days, with 4OHT present for the initial 24 hours to induce recombination **(Figure 4A)**. Control oligodendrocytes extended long, MBP^+^ sheaths that fully wrapped the fibers. In contrast, the MBP signal in cKO cells remained largely restricted to the soma, resulting in significantly shorter and unwrapped microfibers **(Figure 4B-C)**.

**Figure 4.**
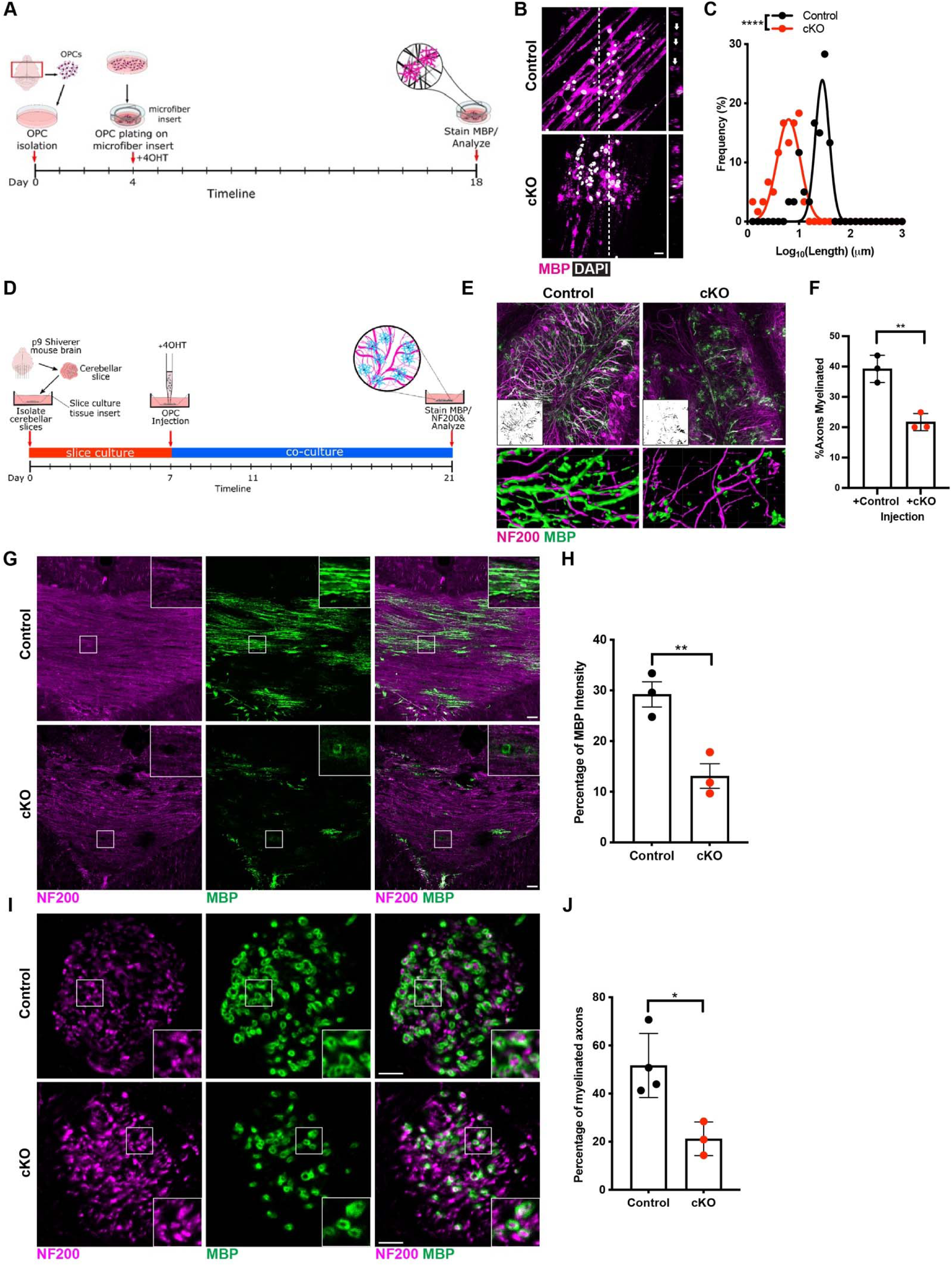
preOL *ADGRG1* is required for myelin formation and repair. **(A)** Schematic of the microfiber assay. **(B)** MBP immunostaining shows ensheathment of microfibers by control and cKO OLs. White arrows in the orthogonal view indicate wrapped microfibers. Scale bar, 10 µm. **(C)** Distribution of sheath lengths in control and cKO OLs, plotted as frequency (%) against log_10_- transformed sheath length (µm). cKO OLs show significantly shorter sheath lengths compared to control cells. 60 total OLs were analyzed per group, and at least 138 sheaths were quantified per condition. Bins were defined with upper log_10_ length limits ranging from 0.1 to 3 in 0.1 increments. n=3 biological replicates. **** p<0.0001. Mann-Whitney test, two-tailed. **(D)** Schematic of co-culture system. **(E)** NF200 (magenta) and MBP (green) staining shows myelination of Shiverer cerebellar slices following addition of control or cKO OPCs. Inserts indicate overlapping NF200 and MBP signals. Bottom panels show 3D reconstructions of myelin sheaths. Scale bar, 50 µm. **(F)** Quantification of the percentage of myelinated axons. Each circle represents one animal. ** p=0.0045. Unpaired two-tailed t-test. **(G)** Representative images of corpus callosum (CC) sections from P14 control and cKO mice stained for NF200 (magenta) and MBP (green). Scale bar, 50 µm. Boxed areas are shown at higher magnification. **(H)** Quantification of MBP coverage in the CC shows a significant reduction in cKO mice. Each circle represents one biological replicate. ** p=0.0098. Unpaired two-tailed t-test. **(I)** Cross-sections of striatal axonal projections from P14 mice stained for NF200 (magenta) and MBP (green). Scale bar, 5 µm. Boxed areas are shown at higher magnification. **(J)** Quantification of the percentage of myelinated axons in control and cKO mice. Each circle represents one biological replicate. * p=0.0163. Unpaired two-tailed t-test. Data are presented as means ± SEM.

We next assessed the myelination capacity of control and cKO oligodendrocytes using an *ex vivo* organotypic slice co-culture system where OPCs were seeded onto cerebellar slices from myelin-deficient Shiverer mice (*29*) **(Figure 4D)**. In the presence of 4OHT, *Adgrg1* will be deleted in the preOLs. Control oligodendrocytes robustly myelinated shiverer axons, while cKO cells failed to form proper myelin sheaths, resulting in a significant reduction in the percentage of myelinated axons **(Figures 4E-F)**. Quantification showed a significant reduction in the percentage of myelinated axons in the cKO-injected slices **(Figure 4F)**.

Finally, we sought to confirm these ensheathment defects *in vivo*. Analysis of the corpus callosum of P14 control and cKO mice revealed defects in myelination in cKO mice. Compared to extensive, longitudinally distributed MBP signals observed along axons in control mice, MBP in cKO mice was dramatically reduced and abnormally accumulated in oligodendrocyte somas **(Figures 4G-H and S3A-B)**. Examination of striatal cross-sections further confirmed a significant decrease in the number of myelinated axons in cKO mice, with the few remaining myelin sheaths appearing incomplete **(Figures 4I-J)**. Taken together, these results demonstrate that *Adgrg1* deletion in preOLs profoundly disrupts axon ensheathment and impairs the formation of compact myelin during development.

### The function of ADGRG1 in axon ensheathment is evolutionarily conserved

To determine if the role of ADGRG1 in axon ensheathment in early development is evolutionally conserved, we performed *in vivo* timelapse imaging of wild-type and *adgrg1/gpr56* mutant embryonic zebrafish spinal cords from 2-3 days post-fertilization (dpf) with a fluorescent marker labeling the OL membrane and F-actin polymers **(Figure 5A)**. Given previous data using this mutant model showing hypomyelination in zebrafish spinal cords (*18*), we hypothesized that zebrafish *adgrg1* regulation of F-actin may also regulate the initial ensheathment of axons. Our results show that *adgrg1^stl13/stl13^*homozygous mutant OLs formed less total length of new ensheathments during the eight-hour timelapse period **(Figure 5B-C)**. The *adgrg1^stl13/+^*heterozygous mutant cells formed an intermediate length of new ensheathments not significantly different from either the wild-type (**Figure 5B-C**) or homozygous mutant conditions. These results suggest that *adgrg1* is necessary for normal length of initial ensheathments in developing zebrafish OLs. Furthermore, the intermediate results of the heterozygous condition suggest a possible gene dosage effect, where reduced levels of *adgrg1* due to haploinsufficiency may cause a disruption to normal ensheathment.

**Figure 5.**
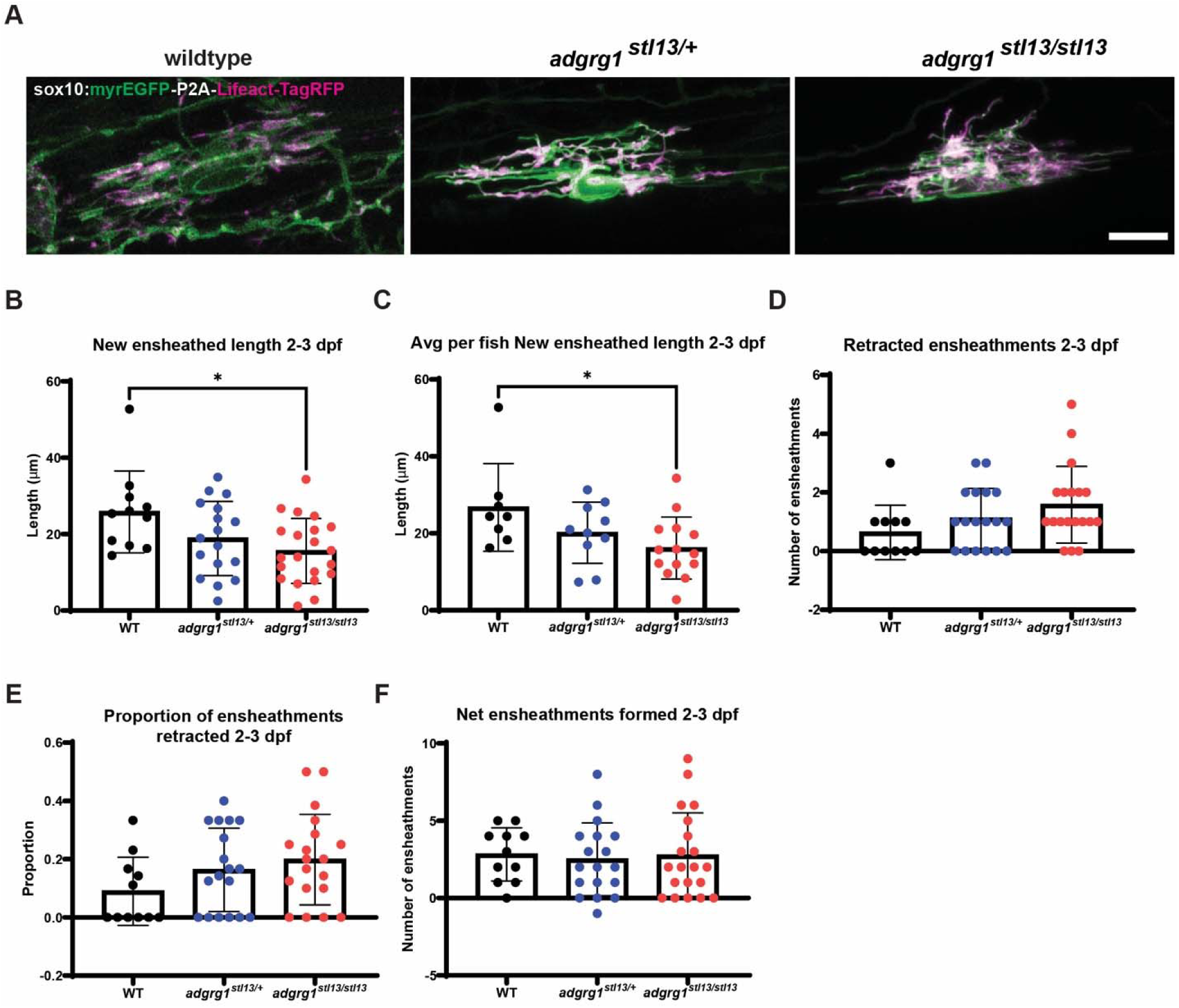
Adgrg1 regulates developmental oligodendrocyte ensheathment of axons in zebrafish. **(A)** Example timelapse images of *sox10:myrEGFP-P2A-Lifeact-TagRFP*-expressing oligodendrocytes in the zebrafish spinal cord at 60 hpf in each genotype. Scale bar, 10 µm. **(B)** Length of new ensheathments formed during timelapse period, each data point represents one cell (* p=0.0201, One-way ANOVA with Tukey’s multiple comparison). **(C)** Each data point represents the average per animal (* p = 0.0412, One-way ANOVA with Tukey’s multiple comparison test WT *vs. adgrg1^stl13/+^* p = 0.2806, WT *vs. adgrg1^stl13/stl13^*p = 0.0319, *adgrg1^stl13/+^ vs*. *adgrg1^stl13/stl13^*p = 0.5359). **(D)** Number of retracted ensheathments during imaging period, each data point represents one cell (ANOVA p = 0.0925). **(E)** Proportion of total ensheathments per cell retracted during imaging period (ANOVA p = 0.1440). **(F)** Net number of ensheathments formed during imaging period, i.e. ensheathments formed minus less ensheathments retracted (ANOVA p =0.9266). WT N = 8 fish, 11 cells, *adgrg1^stl13/+^*N = 10 fish, 17 cells, *adgrg1^stl13/stl13^* N = 14 fish, 21 cells. Data are presented as means ± SEM.

In addition to disruptions in newly ensheathed OL membrane length, our results show a trend towards an increased number of retracted ensheathments by mutant OLs, but it was not significant at the α= 0.05 level (**Figure 5D**). The proportion of retracted ensheathments **(Figure 5E)** and net ensheathments formed **(Figure 5F)** during the imaging period were similar across genotypes. These results suggest that ADGRG1 may aid in stabilization of new sheaths, but its effect on initial ensheathment in zebrafish OLs is mostly related to the elongation of new ensheathments. Together, our results show that Adgrg1 is required for normal initial axon ensheathment by developing OLs in zebrafish embryos.

### ADGRG1 is required in preOLs for axon ensheathment and myelination *in vivo*

To determine how ADGRG1 affects axon ensheathment at the ultrastructural level, we analyzed the optic nerves of control and cKO mice following tamoxifen administration from P4–P7 with booster doses on P10 and P11 **(Figure 6A)**. By P14, transmission electron microscopy (TEM) revealed pronounced defects in axon ensheathment in cKO samples compared with controls. Specifically, cKO optic nerves contained significantly more unmyelinated and partially myelinated axons, whereas control nerves displayed a higher proportion of fully myelinated axons **(Figures 6B–D)**. Moreover, cKO axons exhibited increased outfoldings, the redundant loops of myelin protruding from the main sheath **(Figures 6E–F)**, which is consistent with compromised ensheathment. Notably, the g-ratio (the ratio of axon diameter to the total diameter including myelin) remained unchanged in cKO mice **(Figures 6G–H)**, suggesting that once wrapping commences, myelin thickness is not significantly affected.

**Figure 6.**
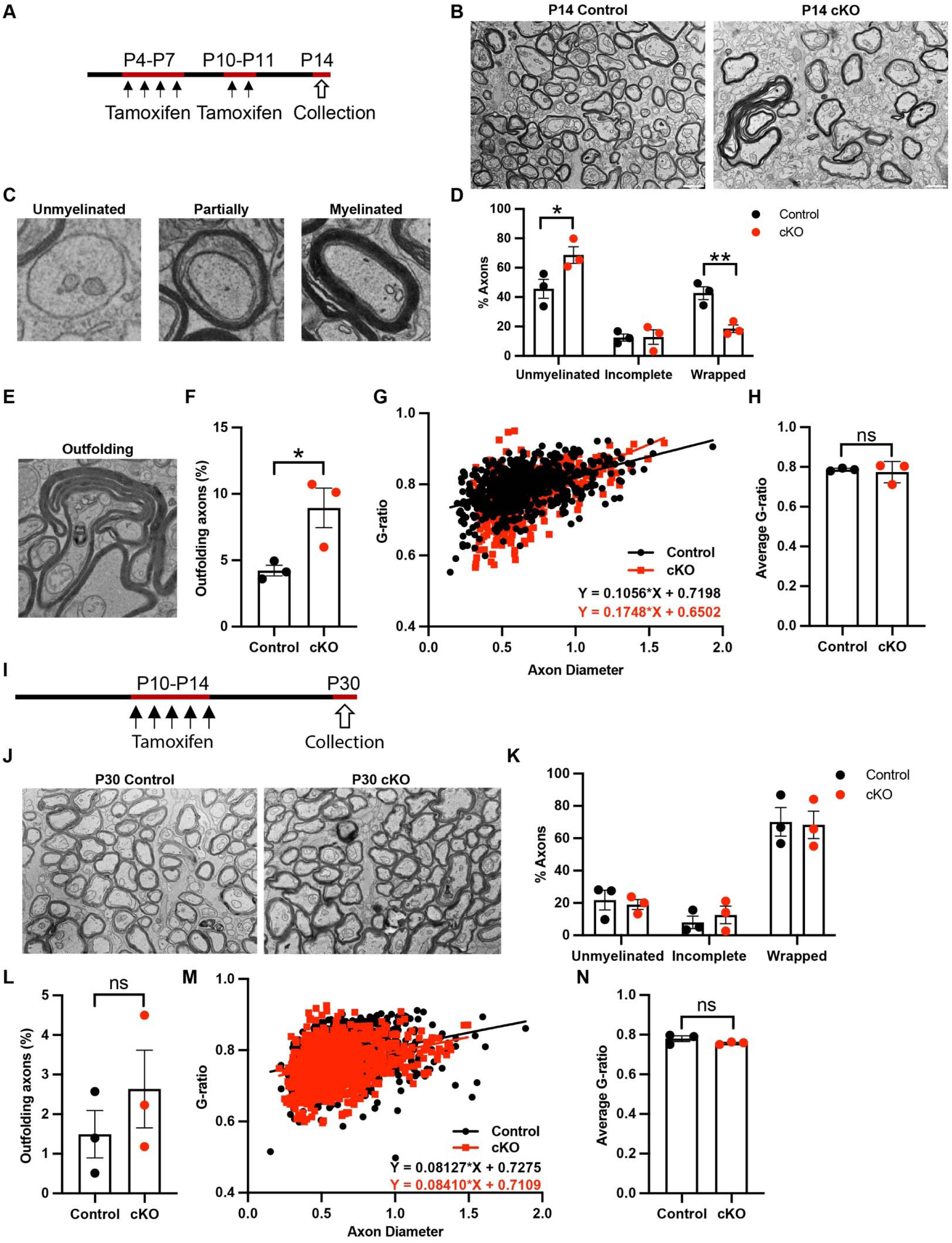
preOL ADGRG1 is required for axon ensheathment *in vivo*. **(A)** Schematic of tamoxifen administration for tissue collection at P14. **(B)** Representative TEM images of optic nerves from P14 control and cKO mice. Scale bar, 1 µm. **(C)** Representative zoomed-in images showing examples of unmyelinated, partially myelinated, and fully myelinated axons. **(D)** Quantification of the percentages of unmyelinated, partially myelinated and myelinated axons. Each circle represents one biological replicate. * p=0.0204, ** p=0.0042. Two-way ANOVA with Bonferroni’s multiple comparisons test. **(E)** Representative TEM image showing axons with myelin outfoldings. **(F)** Quantification of the percentage of outfolding axons in P14 control and cKO mice. Each circle represents one biological replicate. * p=0.0373. Unpaired two-tailed t-test. **(G)** Scatter plot of g-ratio values in optic nerves from control and cKO mice. Over 1100 individual axons were analyzed. Best-fit lines were generated by simple linear regression. n=3 biological replicates. **(H)** Quantification of mean g-ratio values between control and cKO optic nerves at P14. Each circle represents one biological replicate. **(I)** Schematic of tamoxifen administration timeline for tissue collection at P30. **(J)** Representative TEM images optic nerves from control and cKO mice at P30. **(K)** Quantification of unmyelinated, partially myelinated, and myelinated axons at P30. Each circle represents one biological replicate. **(L)** Quantification of axons with myelin outfoldings at P30. Each circle represents one biological replicate. **(M)** Scatter plot of g-ratio values in P30 optic nerves. n=3 biological replicate. **(N)** Quantification of mean g-ratio values between control and cKO optic nerves at P30. Each circle represents one biological replicate. Data are presented as means ± SEM.

To examine whether ADGRG1 is also required for later stages of myelination, we induced *Adgrg1* deletion after the primary ensheathment window (P10–P14) and analyzed optic nerves at P30 **(Figure 6I)**. Under these conditions, no significant differences were observed in unmyelinated, partially myelinated, or fully myelinated axons **(Figures 6J–K)**, nor did we detect obvious changes in outfoldings or G-ratios between cKOs and controls **(Figures 6L–N)**. These data collectively indicate that preOL ADGRG1 is specifically required for the initial process of axon ensheathment but is dispensable once myelin wrapping begins.

## Discussion

In this study, we identify the adhesion GPCR ADGRG1 as a critical, intrinsic regulator of preOL maturation and axon ensheathment. We demonstrate that ADGRG1 is highly enriched in preOLs and that its function within this specific stage is essential for proper myelin formation. Through a combination of *in vitro*, *ex vivo*, and *in vivo* models in both mice and zebrafish, we show that loss of ADGRG1 impairs preOL maturation and prevents them from extending and wrapping myelin sheaths. Mechanistically, we establish that ADGRG1 carries out this function by activating RhoA pathway, providing a direct link between an upstream cell-surface receptor and a key intracellular regulator of myelination **(Figure 7)**.

**Figure 7.**
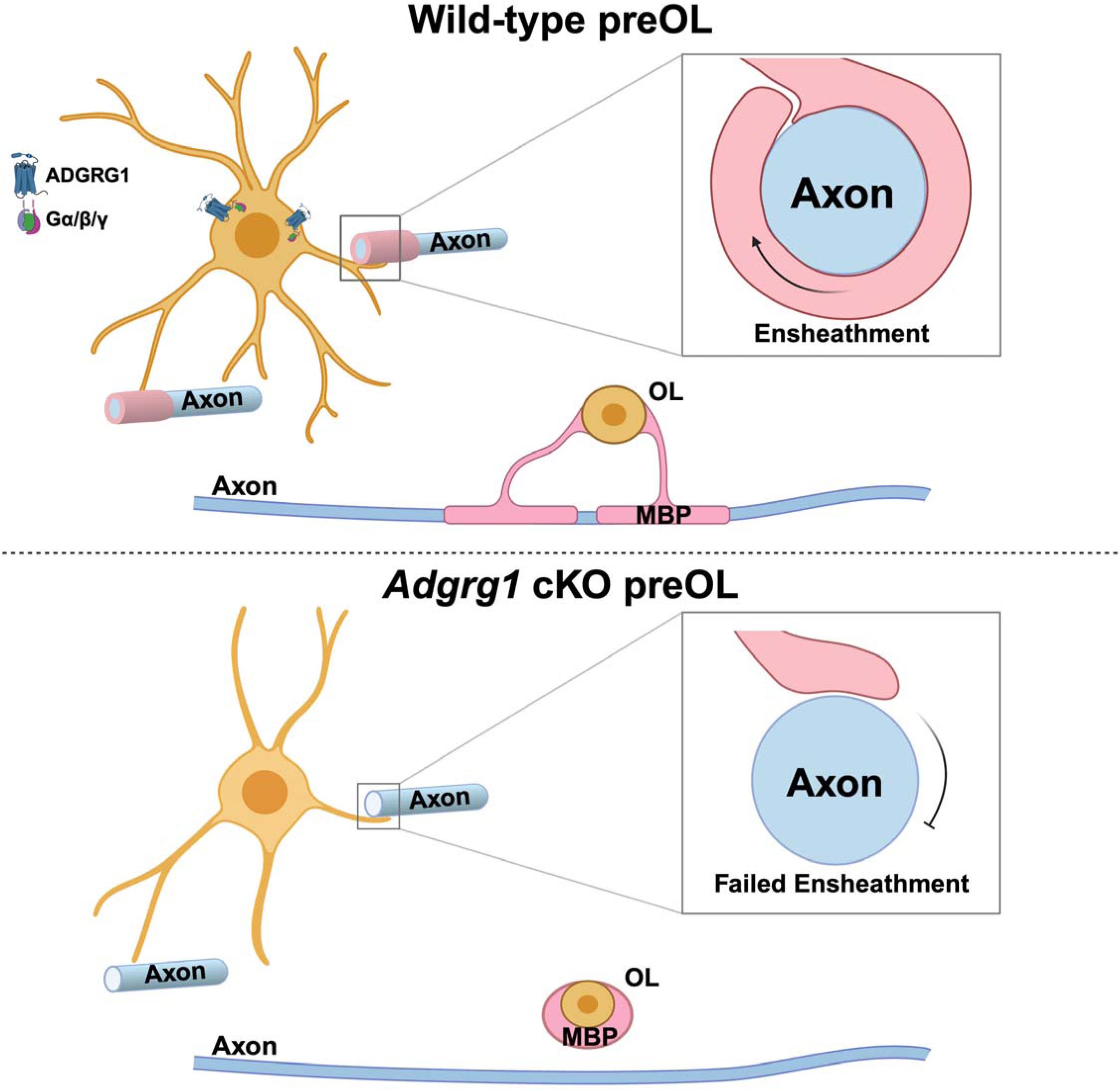
Schematic model of ADGRG1 function in pre-myelinating oligodendrocytes. In wild-type preOLs, ADGRG1 signaling drives morphological maturation, characterized by extensive process branching and distribution of Myelin Basic Protein (MBP) to the processes to initiate axon ensheathment. In contrast, *Adgrg1* conditional knockout preOLs exhibit maturation arrest due to loss of signaling. cKO preOLs display simplified branching morphologies and fail to wrap axons, with MBP aberrantly retained in the cell soma rather than extending into the myelin sheath, resulting in hypomyelination.

Our findings define a novel function for ADGRG1 in preOL maturation that is distinct from its established role in OPC proliferation. Previous studies have shown that global or OPC-specific deletion of *Adgrg1* impairs myelination by reducing the pool of available OPCs through a proliferation defect (*21, 30*). Here, by specifically deleting *Adgrg1* in preOLs, we reveal that ADGRG1 is required for further elaborating preOL processes and axon ensheathment, supporting an indispensable role for ADGRG1 in the later stages of oligodendrocyte lineage development.

The maturation disruption observed in *Adgrg1-*deficient preOLs is mediated by a failure of RhoA activation. Our data show that active, GTP-bound RhoA is present in the processes of control preOLs but is absent in preOLs where *Adgrg1* is conditionally deleted, positioning ADGRG1 as a necessary upstream activator for RhoA signaling at this stage. Ensheathing preOLs have morphologically complex processes. RhoA is an established regulator of cytoskeletal dynamics (*31*) and has previously been shown to act downstream of ADGRG1 signaling (*18*). Accordingly, inhibiting RhoA phenocopied the deficit observed in *Adgrg1* deletion, while activating it was sufficient to rescue the maturation defect in cKO preOLs. Taken together, our study results provide strong evidence that the ADGRG1-mediated RhoA signaling is the primary driver of this process.

RhoA-dependent myelination is less explored compared to other small GTPases like *Rac1* and *Cdc42* (*32*). Here, we show that ADGRG1 is highly expressed in OPCs and preOLs but downregulated in mature oligodendrocytes. This expression pattern parallels the developmental decrease in RhoA activity and supports a model in which high ADGRG1 expression in preOLs drives RhoA-dependent process extension, and its subsequent downregulation in mOLs permits the transition to sheath wrapping, as proposed in the two-step model of myelination (*9*). Our finding is consistent with a recent report by Vale-Silva *et al.*(*19*) that shows a physiological downregulation of GTP-RhoA levels in optic nerves from postnatal day 15 (P15) to adulthood. Together, these findings highlight the importance of temporally regulated ADGRG1 expression in oligodendrocyte lineage for proper myelination.

The conserved function of ADGRG1 in zebrafish highlights its fundamental importance in vertebrate myelination. The failure of oligodendrocytes in *adgrg1* mutant zebrafish to properly extend new sheaths aligns well with the ensheathment defects observed in our mouse models. This evolutionary conservation, combined with ADGRG1’s status as a druggable cell-surface GPCR, makes it a compelling therapeutic target. In many demyelinating diseases, such as multiple sclerosis, remyelination often failed because recruited OPCs differentiate into preOLs that subsequently fail to initiate ensheathment (*33*). Our findings suggest that therapeutically modulating ADGRG1 activity could be a viable strategy to push these arrested cells towards a functional, myelinating state.

While our study establishes a signaling axis from ADGRG1 to RhoA that governs preOL maturation and axon ensheathment, several questions remain. A primary unanswered question is the identity of the endogenous ligand responsible for preOL ADGRG1. In our previous work, we identified microglial Transglutaminase-2 (TG2) as an agonist for ADGRG1 in OPCs (*22*). However, the transition to the preOL stage involves a distinct shift in the function toward physical interaction with axons. It is therefore possible that ADGRG1 relies on a different ligand to trigger the specific RhoA-mediated cytoskeletal reorganization required for ensheathment. This suggests a model where ADGRG1 functionality is temporally modulated by distinct ligand environments as the lineage progress. Furthermore, while our data strongly imply that RhoA acts on the cytoskeleton, direct analysis of F-actin dynamics in preOLs will be necessary. Finally, pharmacological activation of this pathway to rescue myelination defects *in vivo* will be essential for translating these findings toward a therapeutic strategy.

In summary, this study defines a specific and temporally restricted role for ADGRG1 in the oligodendrocyte lineage. We have moved beyond its known function in OPC proliferation to reveal its essential role as a key regulator of preOL maturation and axon ensheathment. By linking ADGRG1 to RhoA activation, we provide a molecular pathway that governs this critical step in the formation of the myelin sheath.

## Materials and Methods

### Animals

All animals were handled according to the guidelines of the Institutional Animal Care and Use Committee at the University of California, San Francisco (IACUC #: AN177440-01A). *Adgrg1^fl/fl^*mice were generated as previously described (*21*). Primers used to check for correct loxP status were: 1) 5’-GGT GAC TTT GGT GTT CTG CAC GAC-3’, 2) 5’-TGG TAG CTA ACC TAC TCC AGG AGC-3’, 3) 5’-CAC GAG ACT AGT GAG ACG TGC TAC-3’. PLP-CreER mice were purchased from The Jackson Laboratory (Strain #: 005975). *Adgrg1^fl/fl^; Plp1-CreERT^+^* mice were crossed with Ai14 mice purchased from The Jackson Laboratory (Strain #: 007914). Primers used to detect the correct Ai14 genotype were: 1) 5’-AAG GGA GCT GCA GTG GAG TA-3’, 2) 5’-CCG AAA ATC TGT GGG AAG TC-3’, 3) 5’-GGC ATT AAA GCA GCG TAT CC-3’, 4) 5’-CTG TTC CTG TAC GGC ATG G-3’. To induce deletion of *Adgrg1* in the myelin ensheathment step, we performed daily intraperitoneal injections into mice with tamoxifen (50 mg/kg, corn oil, Sigma) at postnatal (hereafter referred to as P) day 4-7 for the ensheathment step, and P10-P14 for the myelin wrapping step. Shiverer mice were purchased from The Jackson Laboratory (Strain #:001428) and genotyped using the following primers: 1) 5’-TCC CTG GTG GCA GCT ATG AGC AGA CAC TGA-3’, 2) 5’-CCC CGT GGT AGG AAT ATT ACA TTA CCA GCT-3’, 3) 5’-AGG GGA TGG GGA GTC AGA AGT GAG GAA AGA-3’, 4) 5’-ATG TAT GTG TGT GTG TGC TTA TCT AGT GTA-3’.

### *In vivo* timelapse imaging in zebrafish

All zebrafish work was performed in accordance with institutionally approved protocols at Oregon Health & Science University (OHSU). The zebrafish embryos used in this study were the offspring of heterozygous incrosses of *adgrg1^stl13/+^* zebrafish (*18*). These embryos were genotyped according to the protocol described by Ackerman et al, 2015 (*18*). Zygotes at the one-cell stage were injected with 50 nL of 30 ng/µL *sox10:myrEGFP-P2A-lifeact-TagRFP* DNA construct with Tol2 transposase mRNA. The pTol2-sox10:myrEGFP-P2A-Lifeact-TagRFP plasmid was created by Gateway cloning with pDestTol2PA2, p5E-sox10 promoter, pME-myrEGFP, and p3E-P2A-Lifeact-TagRFP (generated by amplification and restriction digest with XhoI and SpeI of Lifeact-TagRFP from the sox10:Lifeact-TagRFP plasmid (*34*)). Zebrafish embryos were imaged from 60 to 70 hours post-fertilization (hpf) under tricaine anesthesia, with Zeiss LSM-980 airyscan timelapse imaging, with images acquired every 20 minutes. Post-processing in Zen and ImageJ was limited to brightness and contrast.

### Immunohistochemistry

At the time of sacrifice, mice underwent transcardiac perfusion, first with phosphate-buffered saline (1× PBS, pH 7.4), then with 4% paraformaldehyde (PFA, pH 7.4), followed by brain dissection and drop fixation in 4% PFA overnight. The next day, brains were changed to a 30% w/v sucrose solution and allowed to sink. Subsequently, tissue was embedded in OCT, and flash-frozen in a dry ice/100% ethanol bath. The tissue was cryosectioned at 14 µm. For immunostaining, slides were boiled in Retrievagen A solution (ThermoFisher, #BDB550524) for 10 minutes and rinsed three times in 1× PBS. Sections were then incubated with blocking buffer (10% goat serum, 1% bovine serum albumin, and 0.1% Triton X-100 in 1× PBS) followed by primary antibody incubation overnight at 4°C. The following day, sections were washed thrice in 1× PBS, and anti-species secondary antibodies were used at a 1:500 dilution in blocking buffer for 1 hour at room temperature. Slides were then washed and mounted using DAPI Fluoromount-G (VWR, #101442-494).

### Immunocytochemistry

Cells were harvested by 4% PFA fixation for 20 minutes at room temperature. After washing with 1× PBS 3 times, cells were blocked with blocking buffer (10% goat serum, 1% bovine serum albumin, and 0.3% Triton X-100 in 1× PBS) for 1 hour at room temperature, followed by primary antibody incubation overnight at 4°C. The following day, coverslips were washed twice in 1× PBS, and anti-species secondary antibodies were used at a 1:500 dilution in blocking buffer for 1 hour at room temperature. Coverslips were then washed and mounted using DAPI Fluoromount-G (VWR, #101442-494).

### Antibodies and Reagents

For immunohistochemistry studies, we used the following antibodies: rat anti-PDGFRα (BD Biosciences, #562171, 1:250), rat anti-myelin basic protein (MBP, Abcam, #ab7349, 1:200), mouse anti-ADGRG1 (H11 (*21*), 1:200), rabbit anti-BCAS1 (Synaptic Systems, #445003, 1:500), mouse anti-RhoA-GTP (New East Biosciences, #26904, 1:200), mouse anti-Olig2 (Millipore Sigma, #MABN50, 1:200), mouse anti-NF200 (Millipore Sigma, #N0142, 1:500), rabbit anti-RFP (Rockland, #600-401-379, 1:500), and rabbit anti-TCF7L2 (Cell signaling, #2569S, 1:500).

### RNAscope

RNAscope was performed using the RNAscope Multiplex Fluorescent Reagent Kit v2 for fixed cells according to the manufacturer’s instructions. RNAscope Probe-Mm-Tmem2 (ACD, #496041) was used to detect expression of the *Tmem2*.

### Transmission Electron Microscopy

Optic nerves from P14 or P30 mice were fixed by perfusion followed by overnight immersion in a 2% glutaraldehyde/4% paraformaldehyde/0.1 M sodium cacodylate buffer (pH 7.4). Next, optic nerves were postfixed in a 1% osmium tetroxide/1.5% potassium ferrocyanide solution for 1 hour, incubated for 1 hour in 1% aqueous uranyl acetate, and then dehydrated in an increasing gradient of alcohol. Samples were incubated for 1 hour in propylene oxide and then overnight in a 1:1 mixture of propylene oxide and TAAB Epon (Marivac Canada Inc. St. Laurent, Canada). Subsequently, samples were embedded in TAAB Epon and incubated at 60°C for 48 hours. 60 nm ultrathin sections were cut on a microtome, mounted on copper grids, stained with lead citrate, and visualized using a JEOL 1200EX TEM. Images were analyzed using ImageJ to calculate g-ratios and myelinated axons.

### Primary Cell Culture

Primary oligodendrocytes were obtained from *Adgrg1^+/+^*; *Plp-CreER* and *Adgrg1^fl/fl^*; *Plp-CreER* mouse cortices according to the immunopanning protocol (*24*). Briefly, P6-P8 mice cortices were dissected, mechanically homogenized, and digested by papain. Cells were negatively selected against microglia via the use of BSL1 (Vector Laboratories, # L1100) and positively selected for via an anti-PDGFRα antibody (BD Pharmingen, #558774). Oligodendrocyte precursor cells (OPCs) were cultured in a proliferation media containing PDGF-AA (Peprotech, #100-13A) and NT-3 (Peprotech, #450-03) until differentiation was required. To differentiate OPCs, PDGF-AA and NT-3 were removed from the media while T3 (Sigma, #T6397) was added. For the duration of the culture, cells were kept at 37°C and 10% CO_2_, with 50% media changes occurring every 2-3 days.

For immunocytochemistry analyses of the oligodendrocyte lineage, OPCs were plated on 0.01% poly-L-lysine-coated coverslips (Neuvitro, Cat. #: GG-12-pll) in a 24-well format at a density of 10,000 cells. To induce Cre recombination, cultures were treated with 500 nM 4-hydroxytamoxifen (4OHT, Millipore Sigma, #H7904) for the first 24 hours. For pharmacological experiments, compounds were added at the time of differentiation and maintained in the culture media for the duration of the experiment. The following compounds were used: RhoA Activator II (1 µg/mL, Cytoskeleton Inc., Cat. #: CN03-A) and RhoA Inhibitor I (1 µg/mL, Cytoskeleton Inc., Cat. #: CT04-A). Following the experimental period, cells were fixed and processed for immunocytochemistry as described above.

### Cerebellar Slice Culture Preparation

For all cerebellar slice culture experiments, P8-P12 mice were used according to a previously published method (*35*). Briefly, brains and cerebella were extracted from mice and embedded in low-melt agarose blocks, and then sectioned into 300 µm sections in ice-cold Hank’s Buffered Saline Solution (HBSS, Gibco) on a vibratome (Leica, VT1000S). Slices were placed on an insert (Millicell 0.4 µm, Millipore, Cat #: PICM03050) and cultured in slice culture media (SCM) containing 50% MEM, 25% Horse Serum, 25% HBSS, 1× GlutaMax, 5 mg/mL glucose, and 1× Pen-Strep (Fisher Scientific). Slices were allowed 2 days in culture to clear debris and stabilize before any treatment. For immunostaining, the section of insert containing the slices was cut out and moved to a 24-well plate. Next, slices were fixed in 4% paraformaldehyde for 20 minutes, permeabilized in 10% Triton-X 100 for 15 minutes, and stained for MBP and NF200 as described in immunohistochemistry above with the modification that primary antibodies were incubated for 48 hours at 4°C and secondary antibodies incubated for 2 hours at room temperature. During image acquisition, a total of 3 lobes were imaged and averaged for each cerebellar section.

### Cerebellar Co-culture

Shiverer cerebellar slices were obtained in the same manner as described above but were gradually weaned off SCM into serum free media (SFM) containing 50% DMEM, 50% F12, 1% B27, 0.5% N2, 1× GlutaMax, and 1× Pen-Strep. This was done by mixing 2/3 SCM with 1/3 SFM at 3 days *in vitro* and 1/3 SCM with 2/3 SFM at 5 days *in vitro*. Subsequently, only SFM was used. At 7 days *in vitro*, OPCs were microinjected into the Shiverer cerebellar slices. To do this, OPCs were resuspended at a density of 20,000 cells/µL and using a broken glass-pulled Pasteur pipette, 10,000 cells in 0.5 µL were injected per slice. After microinjection, 500 nM 4OHT was added overnight to the media to induce *Adgrg1* knockout. Cultures were incubated for 14 days at 37°C, 5% CO_2_ with SFM media changes every 2-3 days. For experimental conditions, treatments were added at 11 days *in vitro* and kept in the media from that point on. After 14 days of co-culture, sections were treated as described above. For image acquisition, 3 lobes were imaged for each cerebellar section and data averaged. Each section is considered a single biological replicate.

### Microfiber Assay

For the microfiber assay, 12-well fiber inserts containing 2 µm diameter electrically spun fibers were used (AMSBio, Cat. #: TECL-006-1X). Fibers were washed with 20% ethanol and 1× PBS, and coated with 0.01% poly-D-lysine. Subsequently, 30,000 OPCs were plated onto the fibers in each well and the cultures were incubated at 37°C and 10% CO_2_ in differentiation media, with media changes every 2-3 days for a total of 14 days. To induce knockout of *Adgrg1*, 500 nM 4OHT was added overnight to the cell culture media at the time of plating. After 14 days in culture, inserts were stained and mounted (*36*). Briefly, crown inserts were moved to a new 12-well plate and washed thrice with 1× PBS. Next, cells were fixed with 4% paraformaldehyde and washed thrice with 1× PBS. Inserts were stained for MBP overnight at 4°C, incubated with secondary antibodies for 1 hour at room temperature, and mounted using DAPI Fluoromount-G (VWR, #101442-494).

### Confocal Microscopy

All fluorescent sections were viewed and imaged on a Leica TCS SPE confocal microscope and processed using Leica’s LAS X software. 20× images were taken under water immersion, while 40× and 63× magnifications were taken under oil immersion. During image acquisition, all settings such as laser strength, gain, and percent offset were maintained between all experimental groups. 3D reconstruction was performed using Imaris v8.3 (Oxford Instruments).

### RNAseq data analysis

We performed a re-analysis of a previously published single-cell RNA-seq dataset (*20*). Briefly, 5072 cells of the oligodendrocyte developmental lineage from 10 CNS regions of juvenile and adult mouse were analyzed. The clusters expressing synaptic genes and representing vascular and leptomeningeal cell (VLMC) are excluded. Cell-types were identified by unbiased clustering of all cells using Seurat 4.0 (*37*).

### Statistical Analysis

For all studies, images were acquired by blinded investigators. Furthermore, investigators were blinded to genotype throughout the entire study. Blinding was performed by a third party applying coded labels to all subjects, such as slides and genotype samples. Similarly, all quantifications and analyses were performed blinded. Data are presented as mean ± S.E.M. unless otherwise indicated. Asterisks indicate significance: ****p<0.0001, *** p<0.001, **p<0.01, *p<0.05. GraphPad Prism 10 software (GraphPad Software, Inc.) was used for all statistical analyses. Tests used for all analyses are in the figure legends.

## Acknowledgements

The authors would like to thank Dr. Khalida Sabeur for her technical support in OPC immunopanning, Dr. David Bulkley for his assistance with TEM studies, and the Broad Center Microscopy center at the Eli and Edythe Broad Center of Regeneration Medicine. This work was supported in part by the Pediatric Scientist Development Program NICHD grant K12 HD000850 (B.R.) and NINDS grants F31 NS137600 (M.P.), F32NS123005 (C.L.C.), R01 NS094164 (X.P.), R21 NS108312 (X.P.), R01 NS108446 (X.P). The authors declare that they have no conflict of interest.

## Author Contributions

Conceptualization by X.P.; *in vitro* and *in vivo* studies by B.Z., T.L., B.R., A.B.I., B.C., R.S., A.L.T., M.D., C.H.L.; zebrafish studies by M.P., C.L.C supervised by K.R.M; TEM imaging by A.M. supervised by K.R.M.; bioinformatic analysis by A.W., T.L.; and manuscript writing by B.Z., T.L., and X.P.. All authors read and edited the manuscript.

---

**Supplementary Figure 1.**
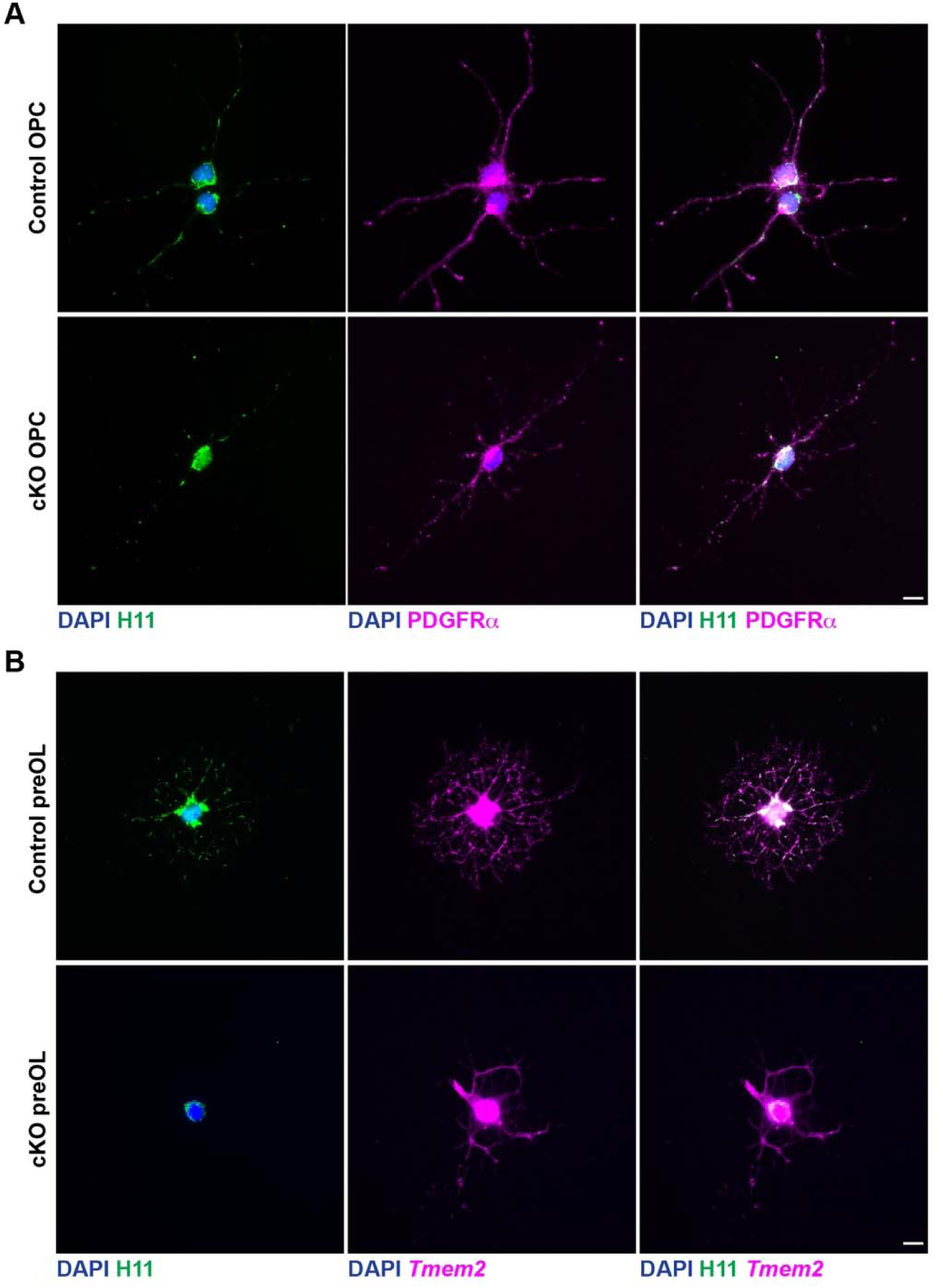
Specific deletion of *Adgrg1* in preOLs but not in OPCs. **(A)** Immunostaining of OPCs immunopanned from control and cKO mice show comparable expression of H11 (green) in PDGFRα^+^ OPCs (magenta), indicating that *Adgrg1* is not deleted at the OPC stage. Scale bar, 10 µm. **(B)** Representative images of preOL staining reveal loss of H11 (green) expression in cKO cells, while control preOL retain H11 signal. *Tmem2* (magenta) labels the preOL stage. Scale bar, 10 µm.

**Supplementary Figure 2.**
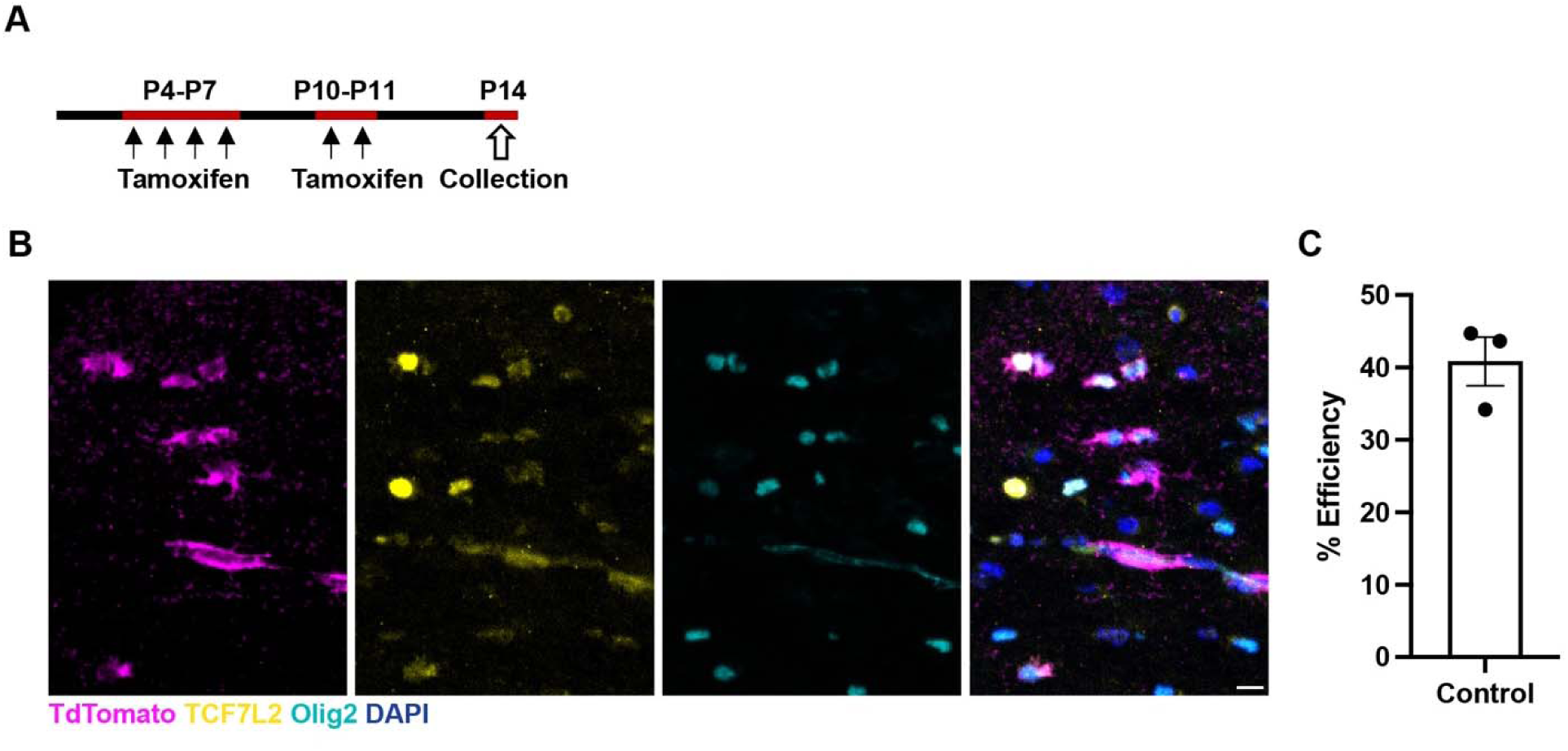
Tamoxifen efficiency in the *Plp^CreERT/+^; Ai14* mice. **(A)** Schematic of the experimental design on tamoxifen-inducible **(B)** Representative immunofluorescence labeling of TCF7L2 (yellow), TdTomato (magenta), Olig2 (cyan) and DAPI (blue) in the corpus callosum of P14 *Adgrg1^fl/fl^; Plp^CreERT/+^; Ai14* mice with tamoxifen induction at P4-P7 and P10-P11. Scale bar, 10 µm. **(C)** Percentage of cells double positive for TCF7L2 and TdTomato, relative to the total number of total TCF7L2^+^ cells. Each circle represents one animal. Data are presented as means ± SEM.

**Supplementary Figure 3.**
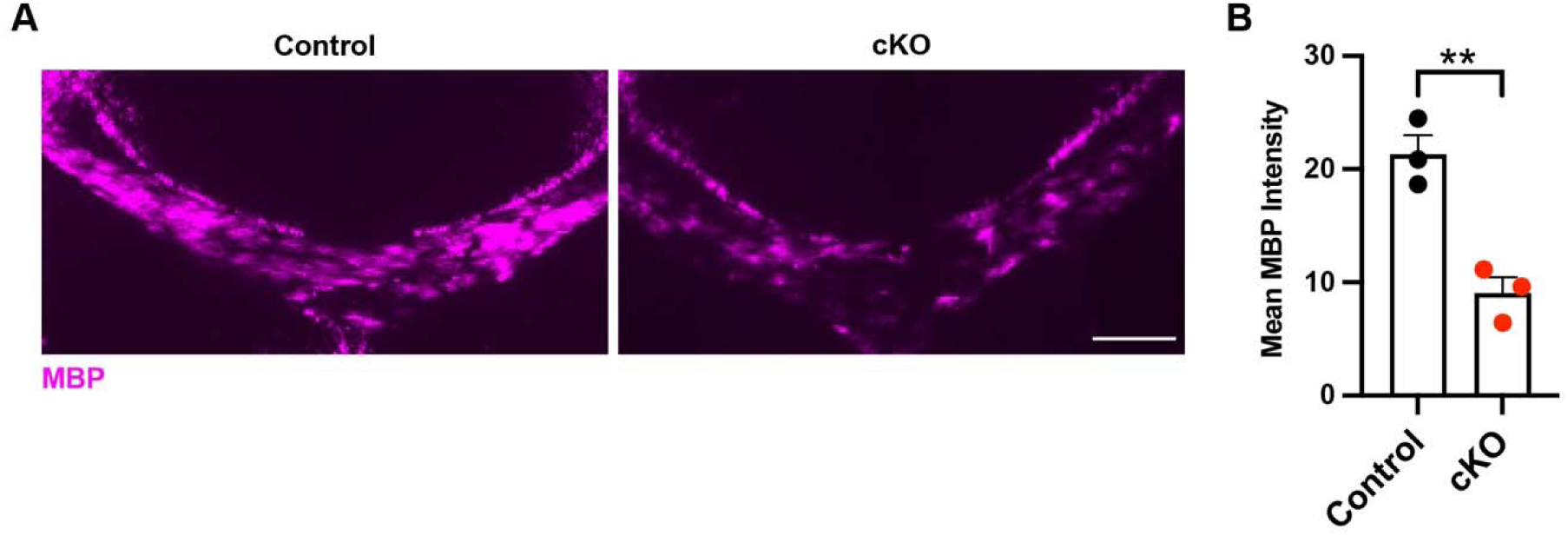
Impaired myelination in the cKO brain at P14. **(A)** Representative coronal sections of the corpus callosum stained for MBP showed reduced myelin signal in cKO mice compared to controls. Scale bar, 50 µm. **(B)** Quantification of mean MBP fluorescence intensity in the corpus callosum showed a significant reduction in cKO mice.

